# Decoding Complex Sounds Using Broadband Population Recordings from Secondary Auditory Cortex of Macaques

**DOI:** 10.1101/614248

**Authors:** Christopher Heelan, Jihun Lee, Ronan O’Shea, David M. Brandman, Wilson Truccolo, Arto V. Nurmikko

## Abstract

Direct electronic communication with sensory areas of the neocortex is a challenging ambition for brain-computer interfaces. Here, we report the first successful neural decoding of English words with high intelligibility from intracortical spike-based neural population activity recorded from the secondary auditory cortex of macaques. We acquired 96-channel full-broadband population recordings using intracortical microelectrode arrays in the rostral and caudal parabelt regions of the superior temporal gyrus (STG). We leveraged a new neural processing toolkit to investigate the choice of decoding algorithm, neural preprocessing, audio representation, channel count, and array location on neural decoding performance. The results illuminated a view of the auditory cortex as a spatially distributed network and a general purpose processor of complex sounds. The presented spike-based machine learning neural decoding approach may further be useful in informing future encoding strategies to deliver direct auditory percepts to the brain as specific patterns of microstimulation.

## Introduction

Electrophysiological mapping by single intracortical electrodes has provided much insight in revealing the functional neuroanatomical areas of the primate auditory cortex. Directly relevant to this work is the role of the macaque secondary auditory cortex in processing complex sounds. Whereas the core of the secondary auditory cortex lies in the lateral sulcus, significant portions of the adjacent belt and parabelt lie on the superior temporal gyrus (STG)^1^. This area is accessible to chronic implantation of microelectrode arrays (MEAs) for large channel-count broadband population recording (see Methods). Earlier seminal research into STG increased our understanding of hierarchical processing of auditory objects by producing meticulous maps of the cellular level characteristics with microwire recordings^1–6^. The implications of non-human primate (NHP) research on human speech processing have been reviewed^7^. Further, a global functional mapping of the NHP auditory system (via tracking metabolic pathways) has been elucidated by fMRI imaging^8^ and 2-deoxyglucose (2-DG) autoradiography^9^.

Other relevant animal studies have examined the primary auditory cortex (A1) in ferrets (guided by single neuron electrophysiological recordings) to unmask the auditory spectrotemporal receptive fields (STRF) underlying tonotopic maps^10, 11^. This work demonstrated encoding properties of the primary sensory neurons in the primary auditory cortex (A1). Studies in marmosets have yielded key insights into audio representation and processing by mapping the STRF beyond A1 into the belt and parabelt regions of the secondary auditory cortex through intrinsic optical imaging^12^. Another recent study examined feedback-dependent vocal control in marmosets^13^ and showed how feedback-sensitive activity of auditory cortex neurons predicts compensatory vocal changes. Importantly, this work also demonstrated how electrical microstimulation of the auditory cortex rapidly evokes similar changes in speech motor control for vocal production (i.e. from perception to action).

Relevant human research has mainly deployed intracranial electrocorticographical (ECoG) surface electrode arrays in STG^14–20^. While surface electrodes are thought to report neural activity over large populations of cells as field potentials, relatively accurate reconstructions of human and artificial sounds have nonetheless been achieved in short-term recordings of able-bodied patients during clinical epilepsy assessment. The recordings generally focus on decoding multichannel low-frequency (0-300 Hz) local field potential (LFP) activity, such as the high-gamma band (70-150 Hz), using linear and nonlinear regression models. LFP decoding has been used to reconstruct intelligible audio directly from brain activity during single trial sound presentations^16, 20^.

Others have implanted ECoG grids and used deep-learning decoding models to reconstruct English language words and sentences^21^. Here, we aimed to access auditory cortical circuits at a much higher spatial and temporal resolution than provided by ECoG or other epicortical electrophysiological techniques. We implanted two 96-channel intracortical arrays in the parabelt areas of the secondary auditory cortex in the rhesus macaque model and successfully decoded multiunit spiking activity to reconstruct intelligible English words and macaque call audio (see Figure 1). Using a novel neural processing toolkit (NPT), we demonstrated the effects of decoding algorithm, neural preprocessing, audio representation, channel count, and array location on decoding performance by evaluating thousands of unique neural decoding models. We showed that high fidelity audio reconstructions of English words and macaque calls can be successfully decoded using multiunit spiking activity.

**Figure 1:**
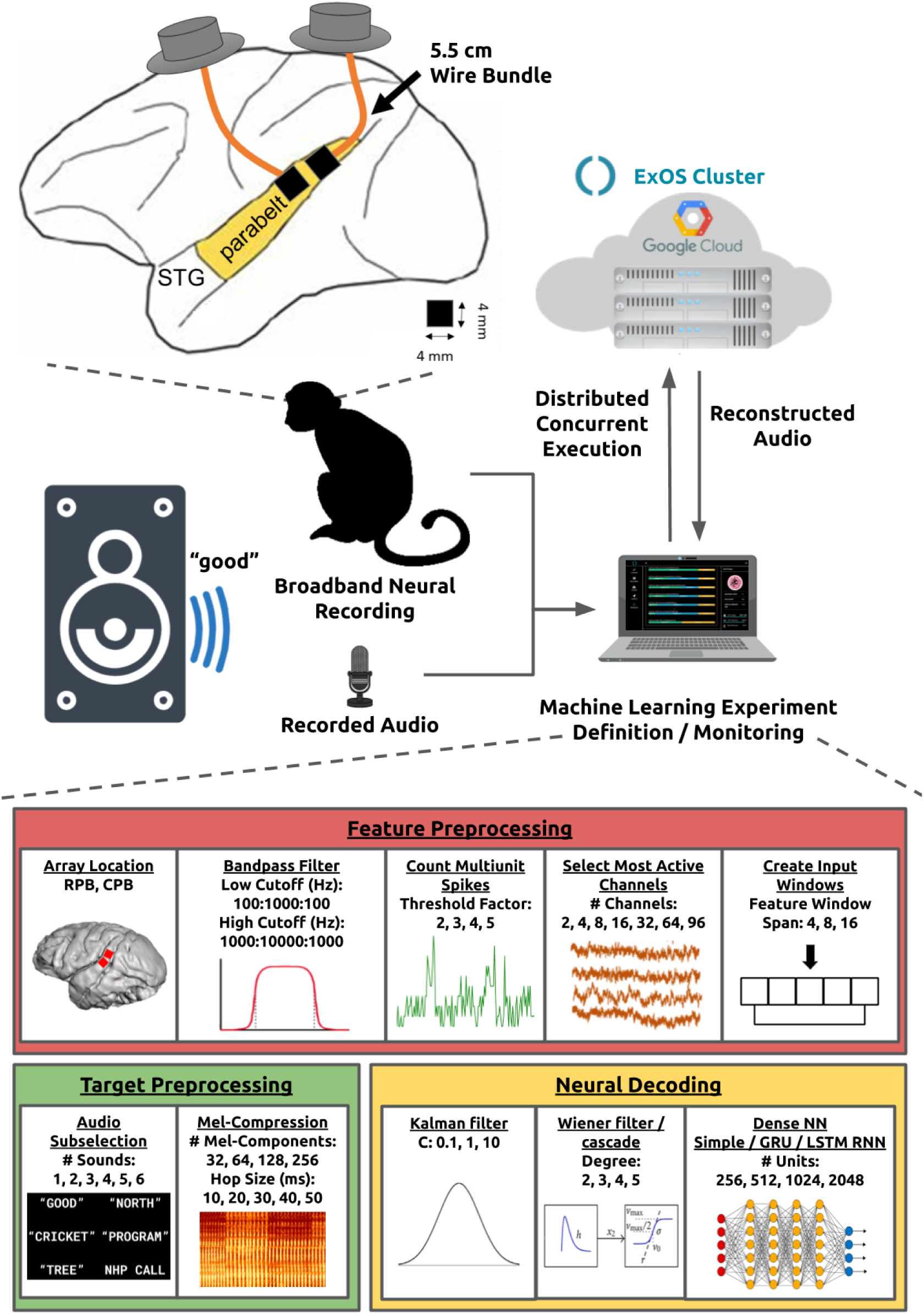
An NHP was implanted with two MEAs in STG. We presented the subject with six recorded sounds and processed neural and audio data on a distributed cluster in the cloud.

## Results

A Summary Video^22^ was prepared to give the reader a concise view of this paper. We used 96-channel intracortical MEAs to wirelessly record^23, 24^ broadband (30 kS/s) neural activity targeting Layer 4 of the STG in an NHP (see Methods). We played audio recordings of 5 English words and a single macaque call over a speaker in a random order. Using a microphone, we recorded the audio playback synchronously with the neural data.

We performed a large-scale neural decoding grid-search to explore the effects of various factors on reconstructing audio from the subject’s neural activity. This grid-search included all steps of the neural decoding pipeline including the audio representation, neural feature extraction, feature/target preprocessing, and neural decoding algorithm. In total, we evaluated 6,059 unique decoding models. Table 1 enumerates the factors evaluated by the grid-search.

**Table 1:** The searched algorithm and hyperparameter space.

| **Neural Decoders** |  |
| --- | --- |
| Kalman filter | C: 0.1, 1, 10 |
| Wiener filter |  |
| Wiener cascade | Degree: 2, 3, 4, 5 |
| Dense Neural Network (NN) | # Units: 256, 512, 1024, 2048 |
| Simple Recurrent NN (RNN) | # Units: 256, 512, 1024, 2048 |
| Gated Recurrent Unit (GRU) RNN | # Units: 256, 512, 1024, 2048 |
| Long Short-Term Memory (LSTM) RNN | # Units: 256, 512, 1024, 2048 |
| **Neural Feature Extraction** |  |
| Array Location: | rostral parabelt (RPB), caudal parabelt (CPB) |
| Filter Low Cutoff (Hz): | 100 to 1000, increments of 100 |
| Filter High Cutoff (Hz): | 1000 to 10000, increments of 1000 |
| Threshold Factor: | 2, 3, 4, 5 |
| # Channels: | 2, 4, 8, 16, 32, 64, 96 |
| Feature Window Span: | 4, 8, 16 |
| **Audio Processing** |  |
| Sounds: | “tree”, “good”, “north”, “cricket”, “program”, macaque call |
| # Sounds: | 1, 2, 3, 4, 5, 6 |
| # Mel-Bands: | 32, 64, 128, 256 |
| Hop Size (ms): | 10, 20, 30, 40, 50 |

### Supplementary Videos

During our analysis, we used Pearson correlation between the target and predicted audio spectrograms as a performance metric for neural decoding models^16^. We present examples in a supplementary Correlation Video^25^ that demonstrates various reconstructions and their corresponding correlation scores. This video aims to provide subjective context to the reader regarding the intelligibility of our experimental results. Additionally, we evaluated neural decoding models with a quantitative intelligibility metric (see Reconstruction Intelligibility). We have also provided a Summary Video that describes the presented findings^22^.

### Neural Decoding Algorithms

Neural decoding models regressed audio targets on neural features. We used the mel-spectrogram^26^ representation of the audio as target variables (see Methods) and multiunit spike counts as neural features. Unless explicitly stated, all models were trained on 5 English words using all 96 neural channels from the RPB array. Both the RPB and CPB array data sets contained 40 repetitions of each word (words randomly interleaved). We reconstructed audio on a trial-by-trial basis (i.e. no averaging across trials).

We evaluated 7 different neural decoding algorithms including the Kalman filter, Wiener filter, Wiener cascade, Dense Neural Network (NN), Simple Recurrent NN (RNN), Gated Recurrent Unit (GRU) RNN, and Long Short-Term Memory (LSTM) RNN (see Methods). We calculated a mean Pearson correlation between the target and predicted mel-spectrogram by calculating the correlation coefficient for each spectrogram component and averaging across components^16^. To mitigate overfitting, we sequentially split the full data set with 80% for training, 10% for validation, and 10% for testing. Results are shown in Figure 2.

**Figure 2:**
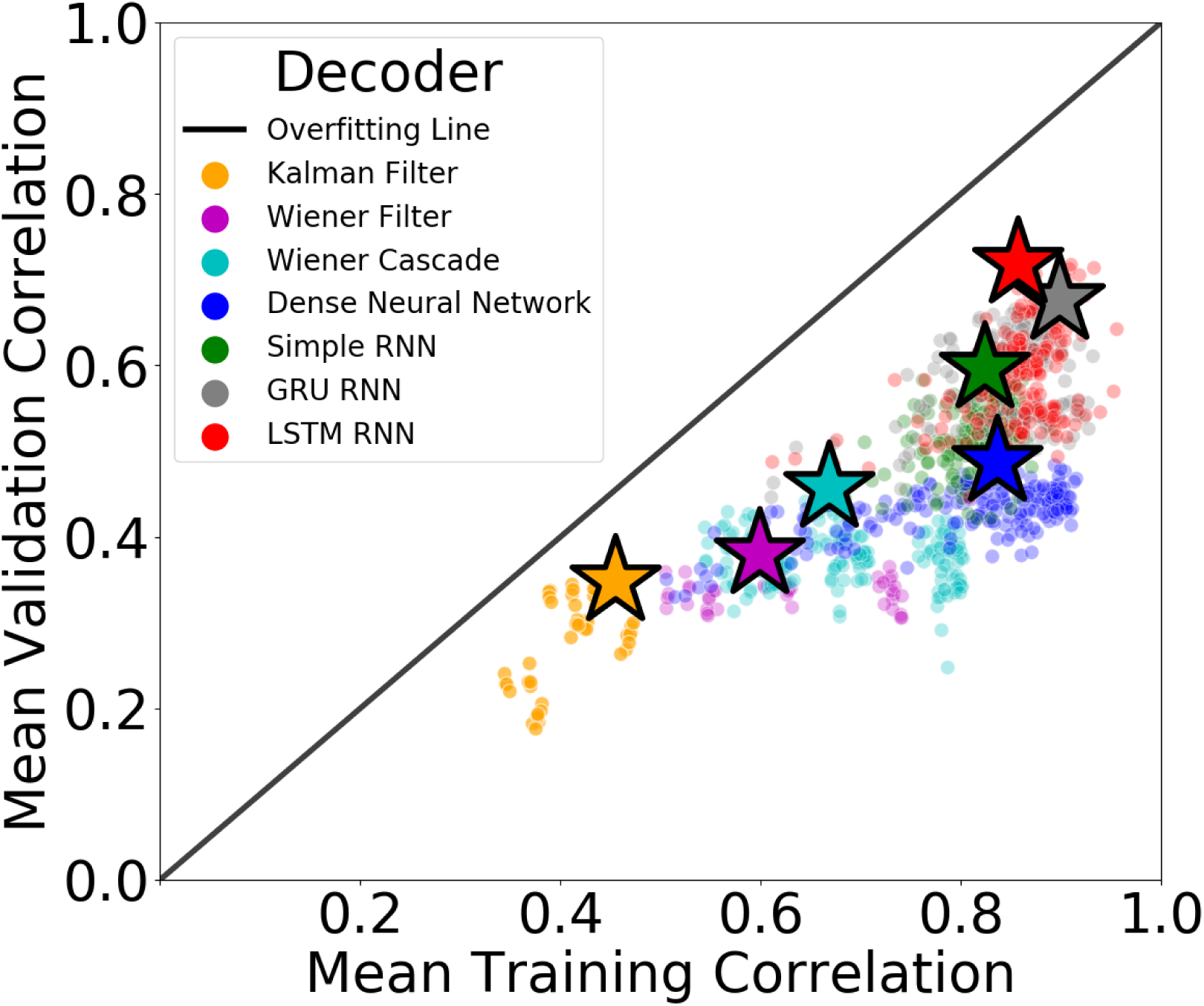
We evaluated the performance of 7 different neural decoding algorithms (color-coded) over both the training data set (x-axis) and validation data set (y-axis). The top performing model for each algorithm is marked with a star. The distance from a given point to the Overfitting Line represents the degree to which the model overfit the training data.

We observed the Kalman filter provided the lowest overall performance with the top model achieving a 0.35 mean validation correlation (MVC) and 0.45 mean training correlation (MTC). The top Wiener filter performed similarly to the Kalman filter on the validation set (0.38 MVC) but showed an increased ability to fit the training set (0.60 MTC). A Wiener cascade of degree 4 beat the top Wiener filter with a MVC and MTC of 0.46 and 0.67, respectively.

A basic densely connected neural network outperformed the top Wiener cascade decoder with a slightly improved MVC (0.49) and MTC (0.84). While the top Simple RNN (0.82 MTC) decoder did not fit the training data as well as the dense neural network, it did generalize better to unseen data (0.60 MVC). Lastly, top GRU RNN (0.68 MVC, 0.90 MTC) and LSTM RNN (0.72 MVC, 0.86, MTC) decoders achieved similar performance with the LSTM RNN providing the best overall performance of all evaluated decoders on the validation set. The LSTM RNN also showed the highest robustness to overfitting with the top model performing only 16% worse on the validation set compared to the training set.

### Neural Feature Extraction

To prepare neural features for decoding models, we bandpass filtered the raw neural data and calculated unsorted multiunit spike counts across all channels.

#### Bandpass Filter

We first bandpass filtered the raw 30 kS/s neural data using a 2nd-order elliptic filter in preparation for threshold-based multiunit spike extraction. To explore the effect of the filter’s low and high cutoff frequencies, we performed a grid-search that evaluated 99 different bandpass filters. We found that using a low cutoff of 500-600 Hz and a high cutoff of 2000-3000 Hz provided a marginal improvement in decoding performance (see Figure 3).

**Figure 3:**
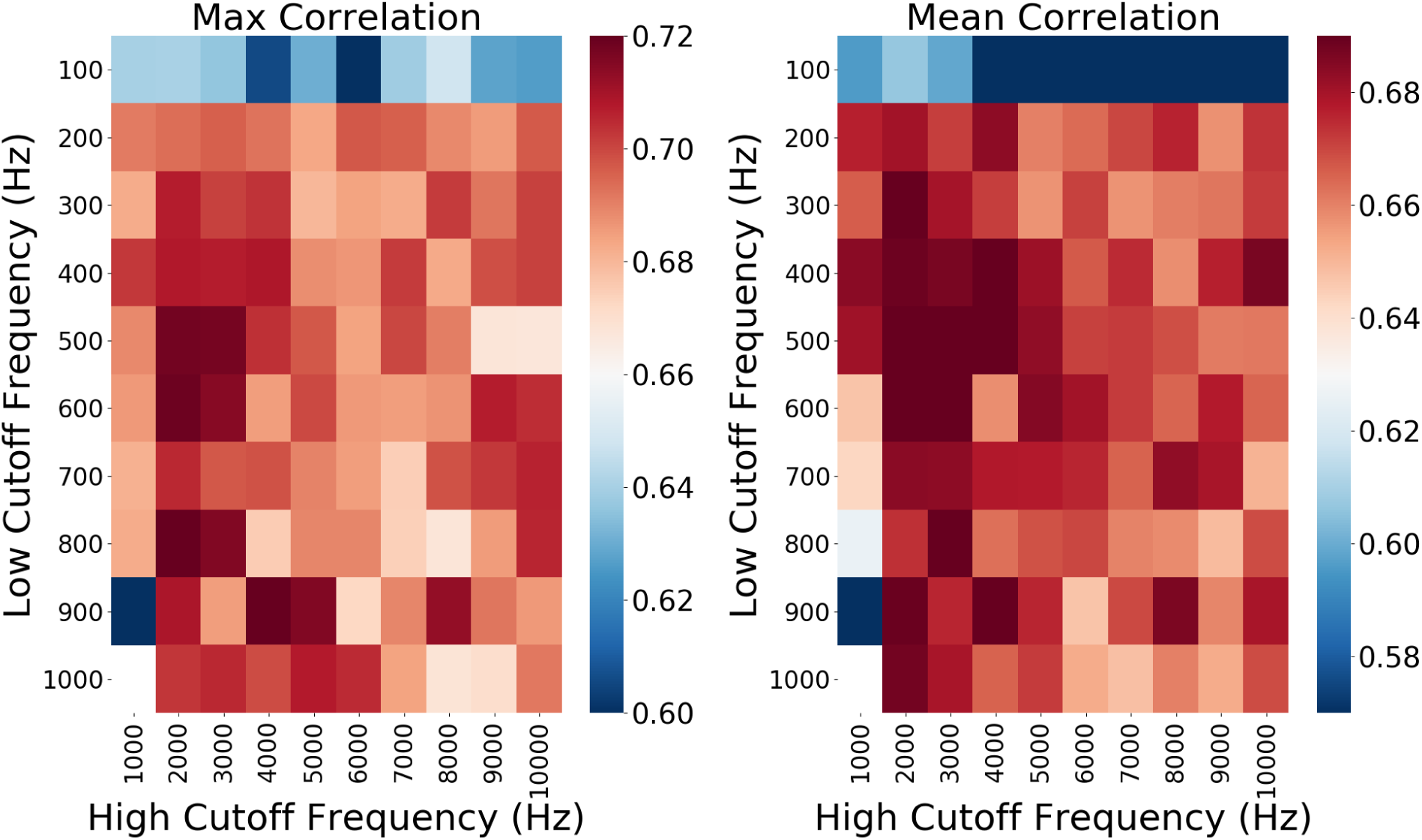
A performance heat-map of the grid-searched bandpass filter cutoff frequencies. We evaluated 99 different bandpass filters. The left and right plots show max and mean performance, respectively, for the filters. We observed a marginal improvement in decoding performance when using a low cutoff of 500-600 Hz and a high cutoff of 2000-3000 Hz.

#### Multiunit Spike Counts

We used multiunit (i.e. unsorted) spike counts as neural features for decoding models. After filtering, we calculated a noise level for each neural channel over the training set using median absolute deviation^27^. These noise levels were multiplied by a scalar threshold factor to set an independent spike threshold for every channel (same threshold factor used for all channels). We extracted negative threshold crossings from the filtered neural data (i.e. multiunit spikes) and binned them into spike counts over non-overlapping windows corresponding to the audio target sampling (see Audio Representation).

As shown in Figure 4, while we observed optimal values for the threshold factor, we found no clear pattern across decoding algorithms. The Wiener cascade was the most sensitive to thresh-old factor with a 16% difference between the most and least optimal values. LSTM RNN decoders were the least sensitive to threshold factor with a 2% difference between the best and worst performing values.

**Figure 4:**
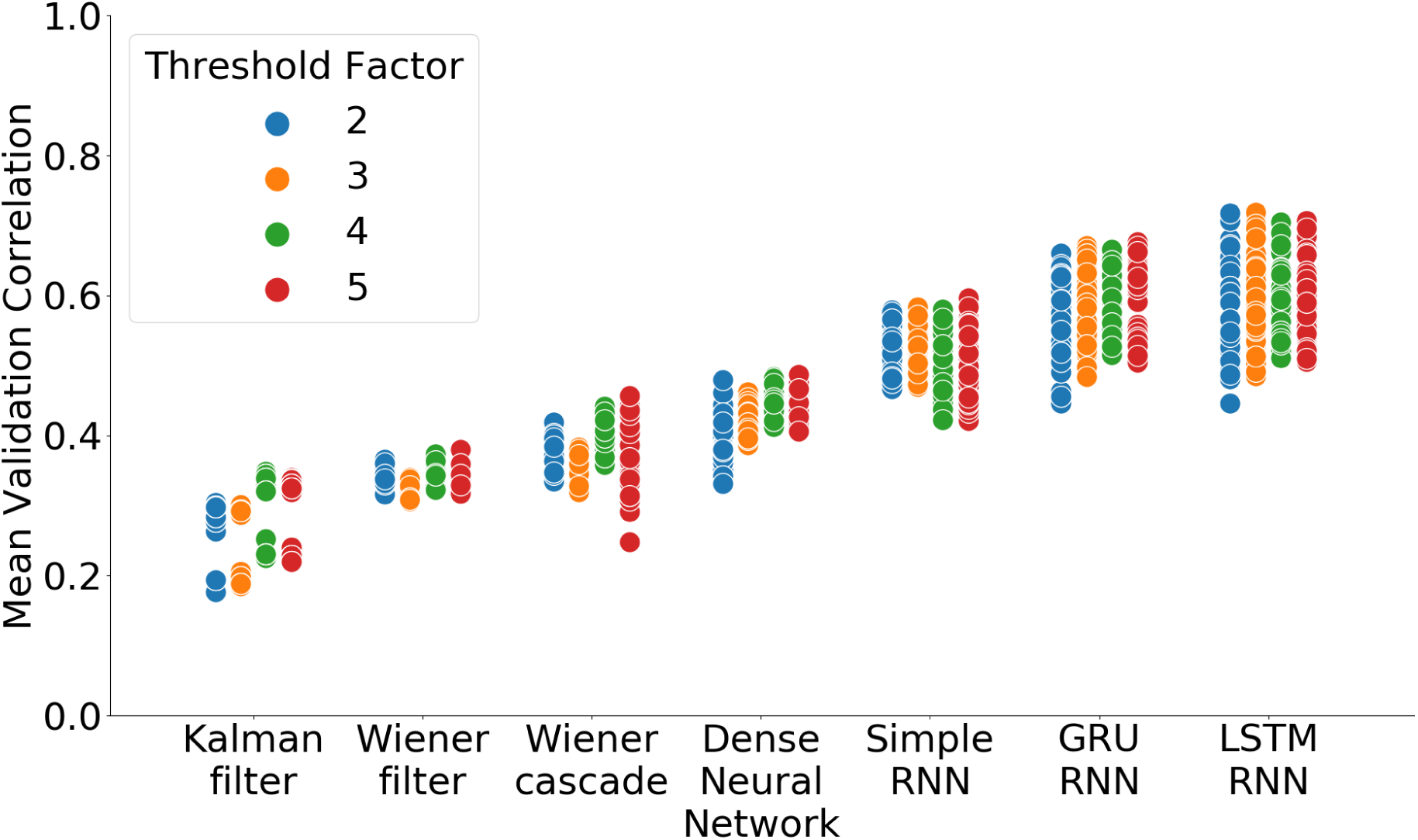
Each marker represents a unique neural decoding model (different hyperparameters) for a given decoding algorithm and threshold factor. Threshold factor did not show a consistent effect on performance across decoding algorithms.

#### Feature Window Span

Except for the Kalman filter which processed a single input at a time, we used a window of sequential values for each feature as inputs to the decoding models. Each window was centered on the current prediction with its size controlled by a “feature window span” hyperparameter (length of the window not including the current value). We evaluated 3 different feature window spans (4, 8, and 16) as shown in Figure 5.

**Figure 5:**
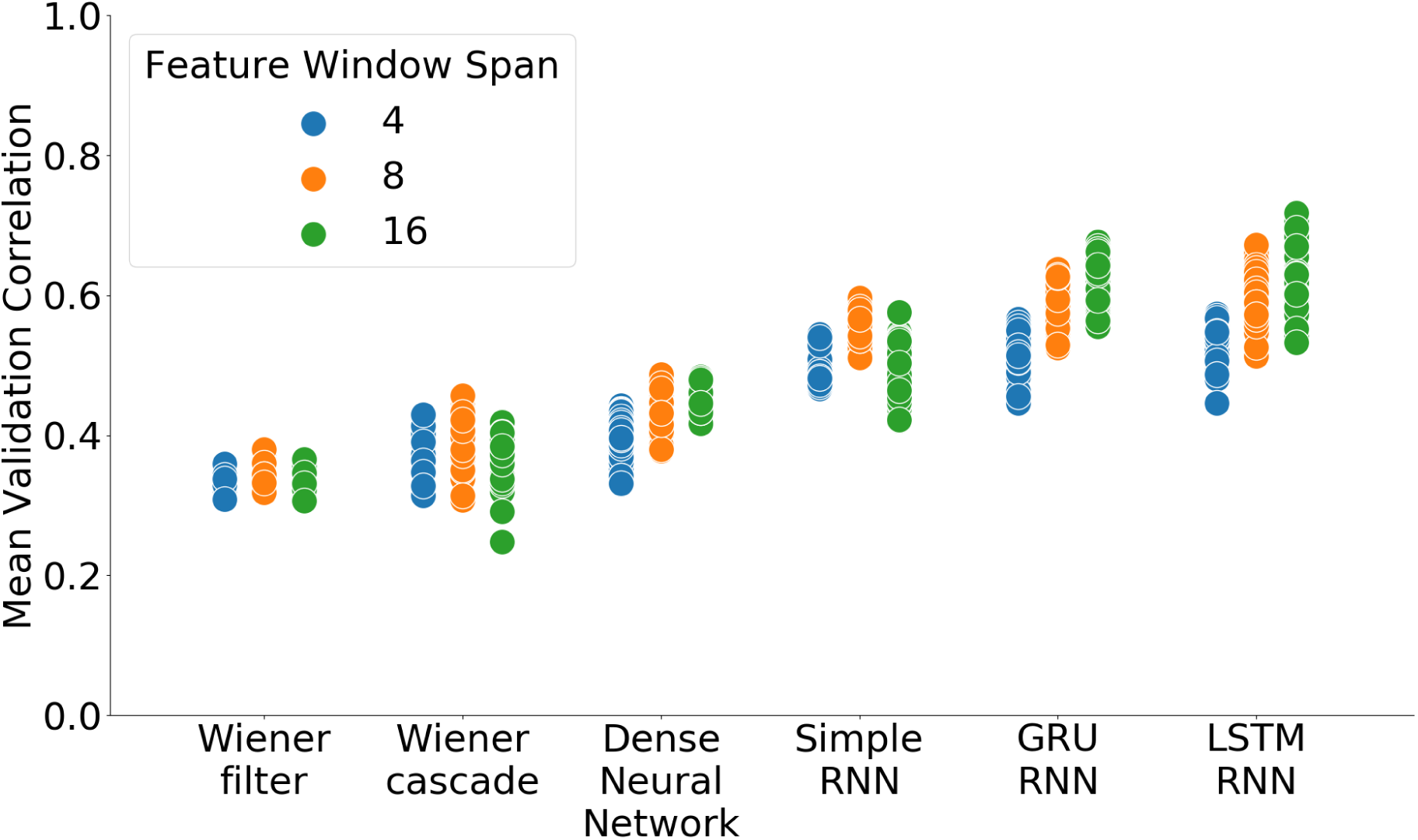
Each marker represents a unique neural decoding model (different hyperparameters) for a given decoding algorithm and feature window span. The GRU RNN and LSTM RNN showed higher performance with a larger feature window span (16). A shorter feature window span (8) achieved top performance for the other neural decoding algorithms.

Both the GRU RNN and LSTM RNN showed higher performance with a larger feature window span (16); however, the other decoding algorithms achieved top performance with a smaller value (8). The LSTM RNN showed the most sensitivity to this hyperparameter with a 20% difference between a feature window span of 16 and 4. The Wiener filter was the least sensitive with a 5% difference between the best and worst performing values. These results demonstrated the importance of determining the optimal feature window span when leveraging LSTM RNN neural decoding models.

#### Channel Count

To investigate the effect of channel count on decoding performance, we ordered neural channels according to highest neural activity (i.e. highest counts of threshold crossings) over the training data set. For 7 different channel counts (2, 4, 8, 16, 32, 64, 96), we selected the top most active channels and built models using only the subselected channels.

As shown in Figure 6, we observed improvements in performance as channel count was increased. We found that selecting the 64 most active neural channels achieved the best performance on the validation set for an audio data set consisting of 5 English words.

**Figure 6:**
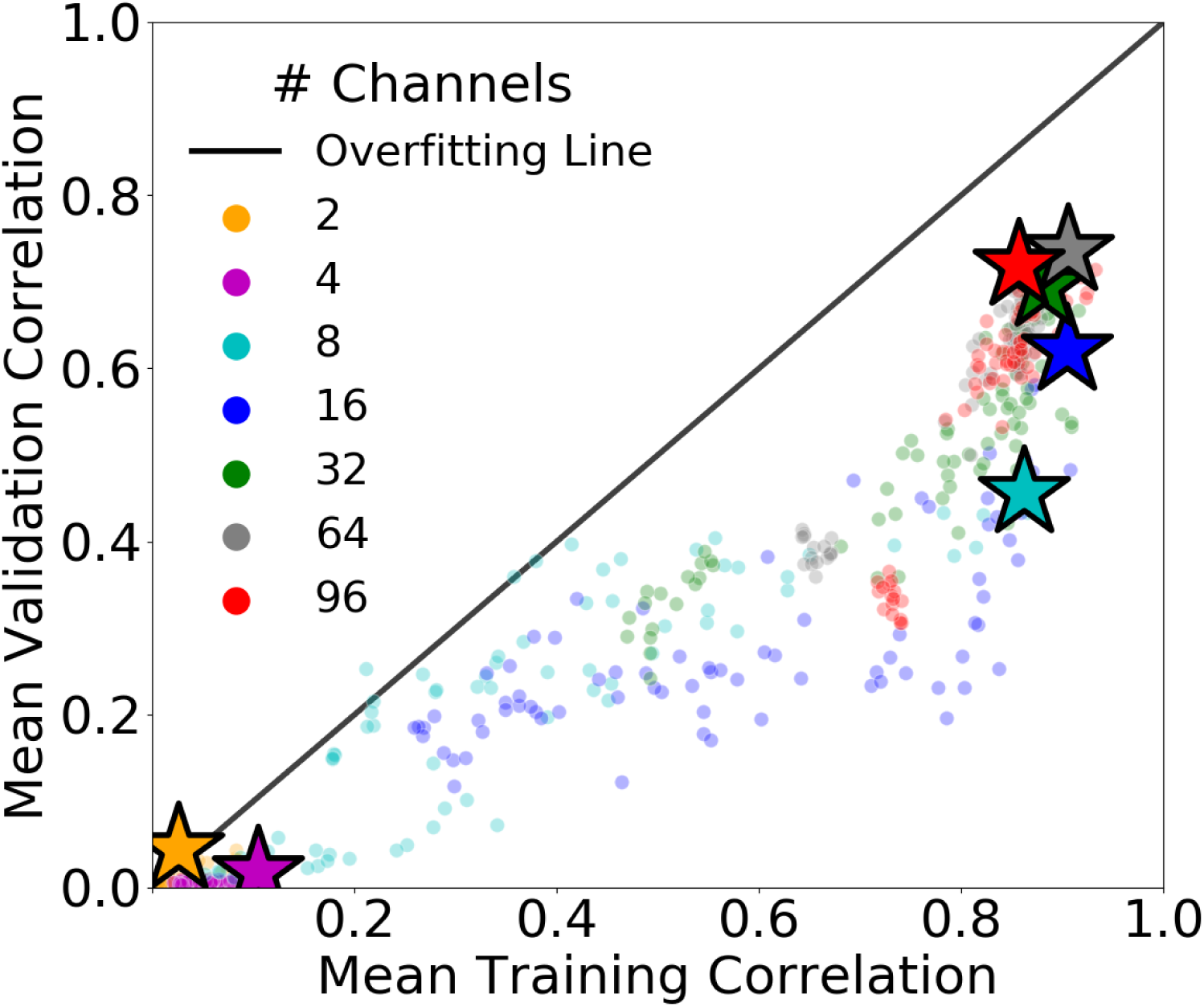
Selecting the 64 most active neural channels provided the best performance on the validation set. The top performing model for each channel count is marked with a star. The distance of a given point to the Overfitting Line represents the degree to which a model overfit. All models shown here were LSTM RNN neural decoders trained on an audio data set of 5 English words.

### Audio Representation

#### Mel-Compression

We converted raw recorded audio to a mel-frequency spectrogram^26^. We examined two hyperparameters for this process including the number of mel-bands in the representation and the hop size for the short-time fourier transform (STFT). We evaluated 4 different values for the number of mel-bands (32, 64, 128, and 256) and 5 different hop size values (10ms, 20ms, 30ms, 40ms, and 50ms). To reconstruct audio, the mel-spectrogram was first inverted, and the Griffin-Lim algorithm^28^ was used to reconstruct phase information.

During preliminary studies, we observed increased decoder performance (mean validation correlations) when using fewer mel-bands and longer hop sizes. However, the process of performing mel-compression with these hyperparameter settings caused even perfectly reconstructed mel-spectrograms to become unintelligible. Therefore, by subjectively listening to the audio reconstructions, we determined that a hop size of 40ms and 128 mel-bands provided the best trade-off of decoder performance and result intelligibility.

#### Audio Complexity

To investigate the effect of audio complexity on decoding performance, we subselected different sets of sounds from the recorded audio data prior to building decoding models. Audio data sets containing 1 through 5 sounds were evaluated using the following English words: “tree”, “good”, “north”, “cricket”, and “program” (for macaque call results, see Macaque Call Reconstruction). For each audio data set, we then varied the channel count (see Neural Feature Extraction) across 7 different values (2, 4, 8, 16, 32, 64, 96).

As shown in Figure 7, we found the optimal channel count varied with task complexity as increased channel counts generally improved performance on more complex audio data sets. However, we also observed decoders overfit the training data when the number of neural channels was too high relative to the audio complexity (i.e. number of sounds). Similar results have been observed when decoding high-dimensional arm movements using broadband population recordings of the primary motor cortex in NHPs^29^.

**Figure 7:**
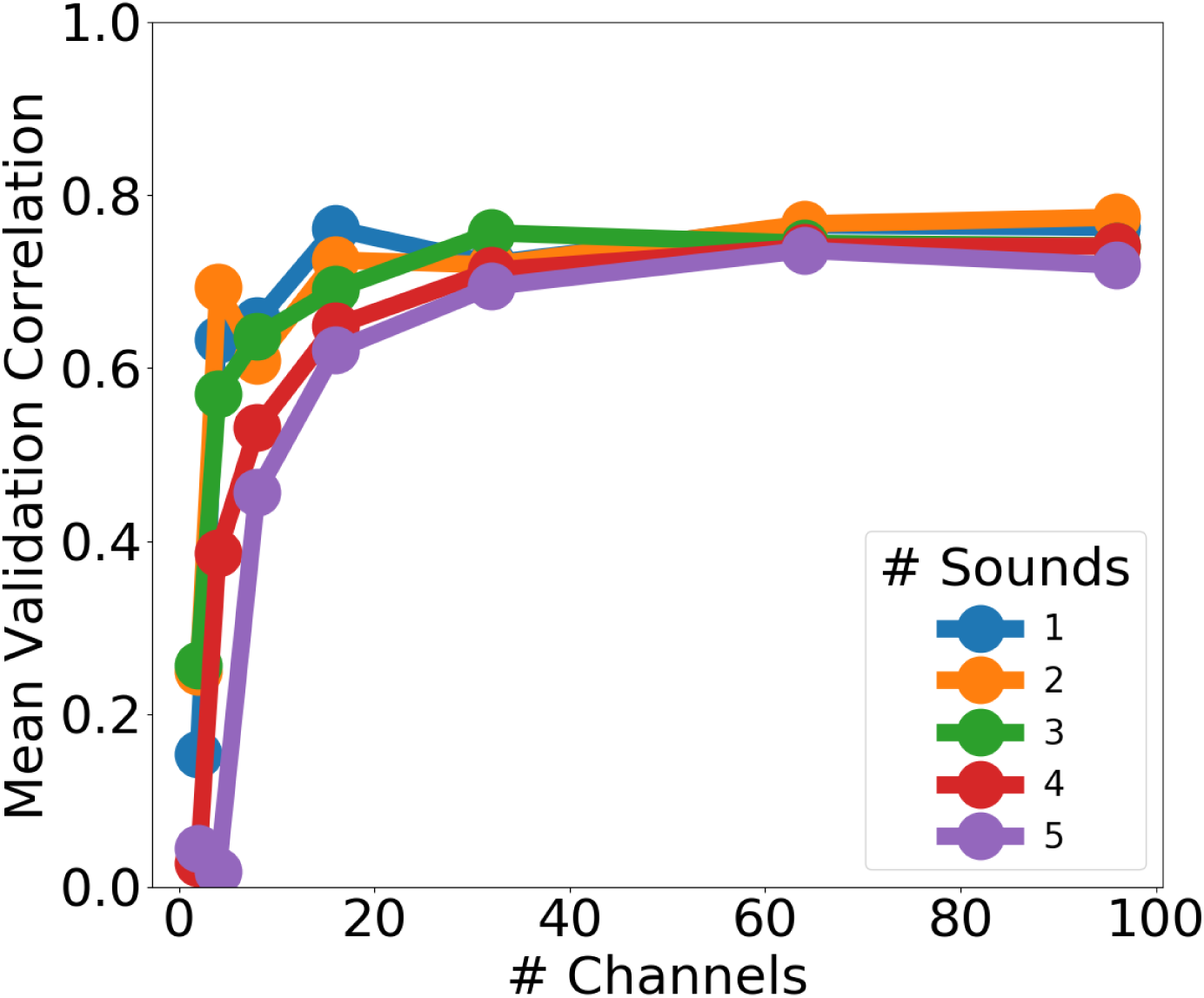
In general, increased channel counts improved performance on more complex audio data sets.

### Array Location

We implanted an NHP with two 96-channel MEAs in the STG, one array in the rostral parabelt (RPB) and a second array in the caudal parabelt (CPB) (see Figure 1). We repeated experiments for each array and compared decoding performance in the two different STG locations (see Figure 8). We successfully reconstructed intelligible audio from both arrays with the RPB array slightly outperforming the CPB array. The similarities between the rostral and caudal results suggest the neural representations of complex sounds are spatially distributed in the STG network. Future work will explore the benefit of synchronously recording from both arrays to enable decoding of more complex audio data sets.

**Figure 8:**
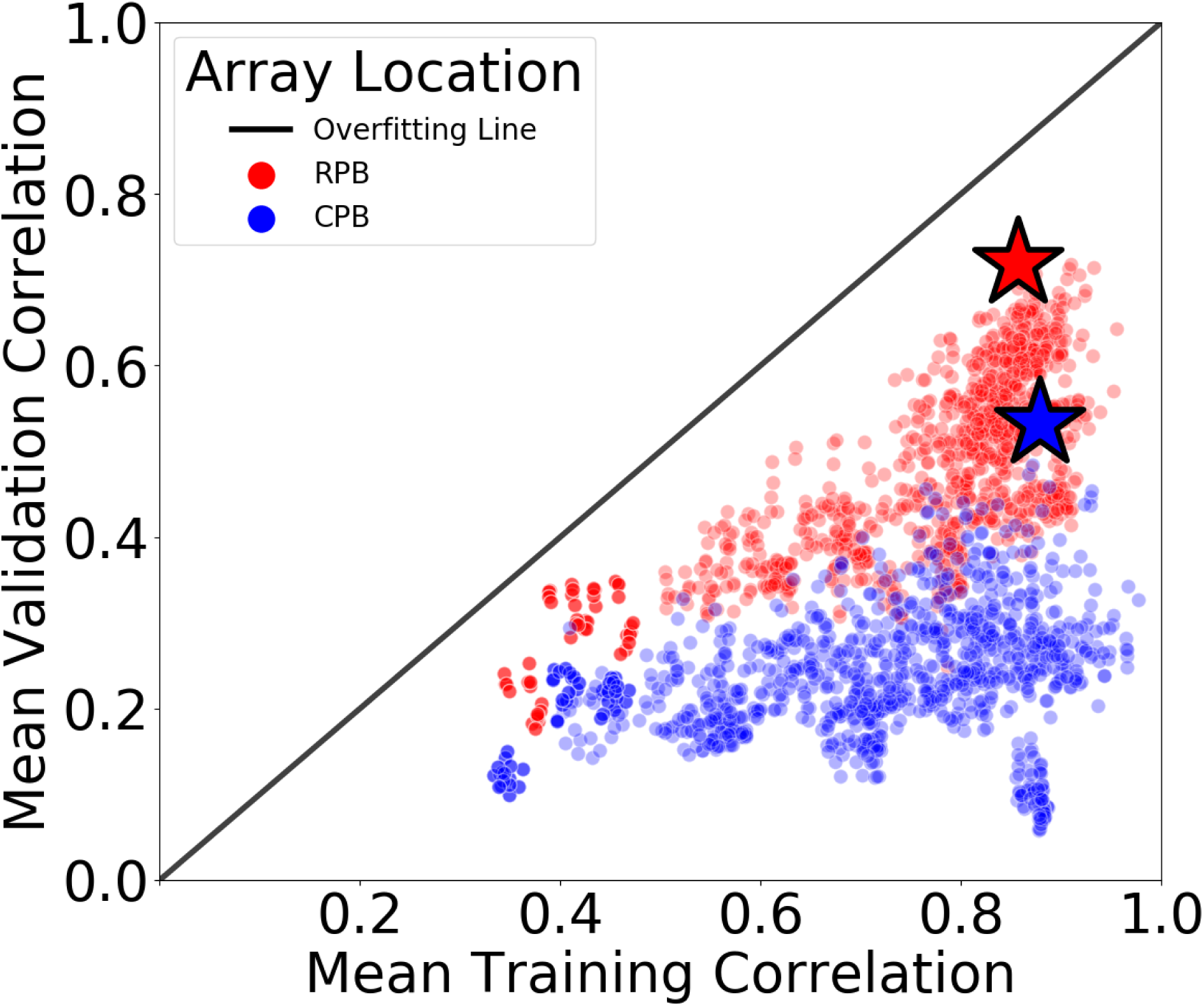
We generated intelligible audio reconstructions using both the RPB and CPB arrays with RPB models achieving higher performance than CPB models on the validation set.

### Reconstruction Intelligibility

The Extended Short-Time Objective Intelligibility (ESTOI) algorithm estimates the average intelligibility of noisy audio samples across a group of normal-hearing listeners^30^. Others have previously utilized ESTOI as a means of quantifying the intelligibility of audio reconstructions decoded from epicortically recorded STG neural activity^21^. We calculated ESTOI scores for all decoding models to analyze the effect of neural decoding algorithm and channel count on reconstruction intelligibility (see Figure 9).

**Figure 9:**
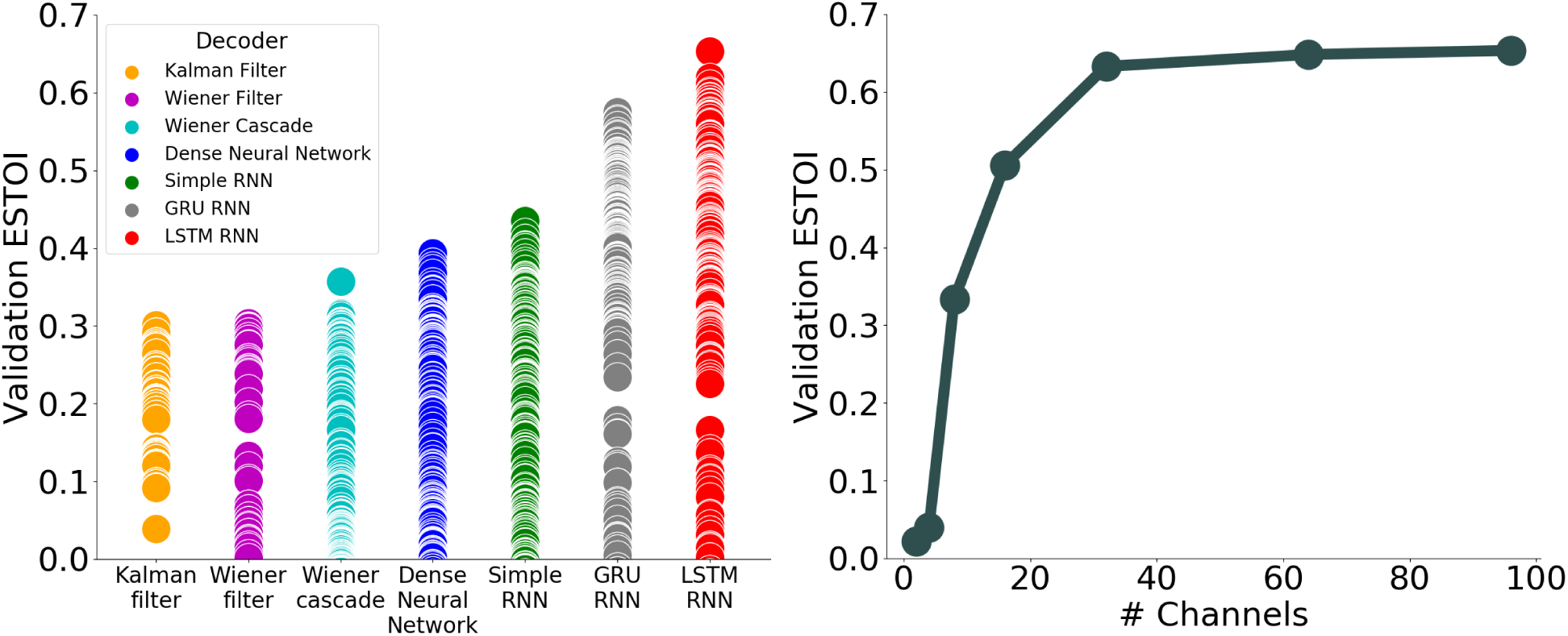
Each marker represents a unique neural decoding model (different hyperparameters) for a given decoding algorithm. LSTM RNN decoding models created reconstructions with the highest intelligibility scores. Intelligibility improved as channel count increased with performance asymptoting at 32 channels.

We observed similar trends in ESTOI performance to those shown in Figure 2 and Figure 7. LSTM RNN neural decoders produced the most intelligible audio reconstructions, and decoding performance saturated using 32 channels of neural activity. The top performing neural decoding model achieved an ESTOI of 0.65 on the validation set.

### Macaque Call Reconstruction

In addition to the English words enumerated in Table 1, we investigated neural decoding models that learned to reconstruct macaque call audio from neural activity. One audio clip of a macaque call was randomly mixed in with the English word audio during the passive listening task.

The addition of the macaque call to the audio data set improved the achieved mean validation correlation from a 0.72 to a 0.87 despite increasing target audio complexity (see Figure 10). This was due to the macaque call containing significant higher frequency spectrogram components compared to audio of the 5 English words (see Figure 12). By successfully learning to predict these higher frequency spectrogram bands, neural decoding models achieved a higher average correlation across the full spectrogram than when those same bands contained mostly noise. While adding the macaque call improved the average correlation scores, it also decreased the top ESTOI score from a 0.65 to a 0.63.

**Figure 10:**
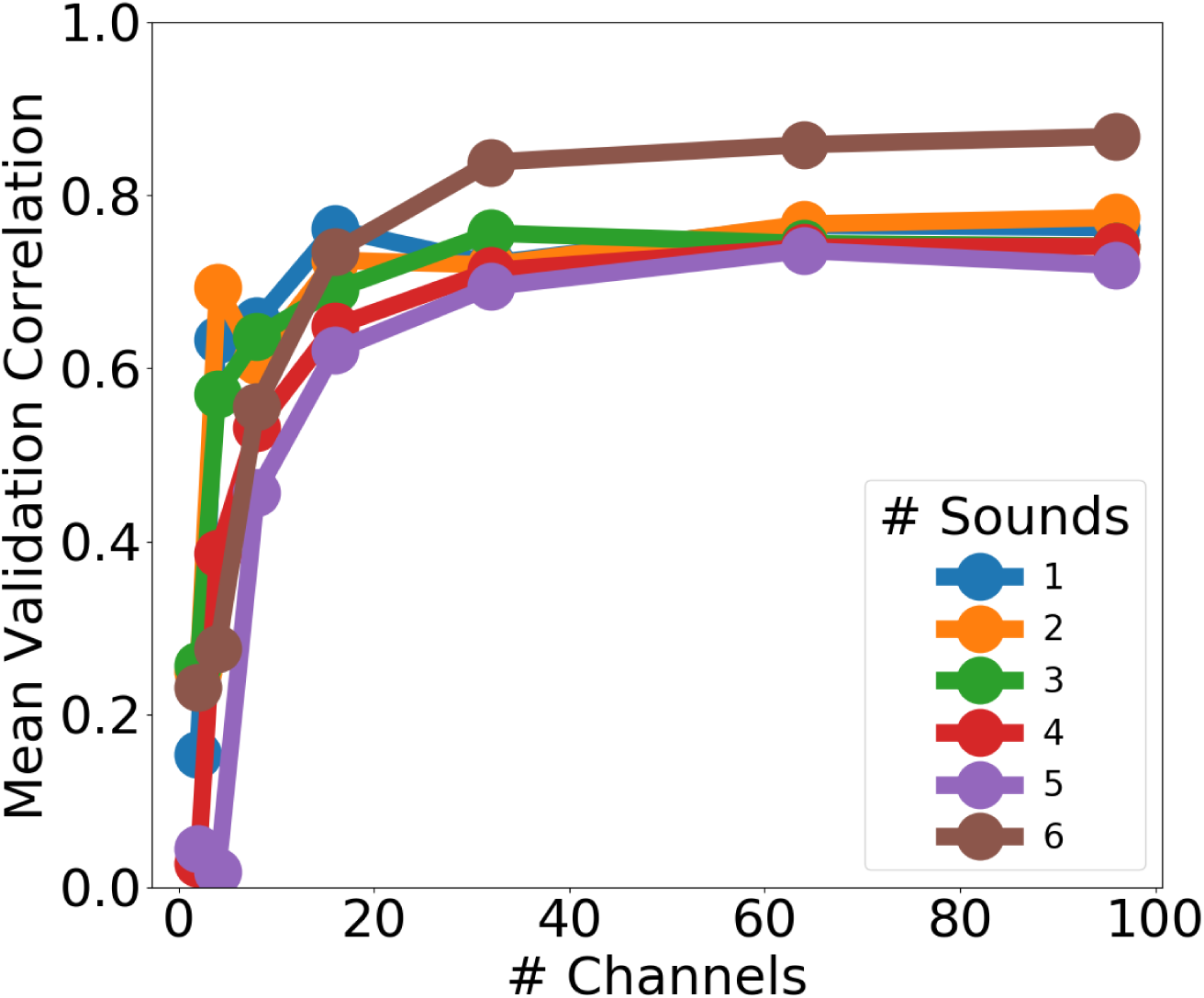
The addition of macaque call audio to the 5 English words improved average correlation scores across the audio spectrogram on the validation set. Performance continued to asymptote at 32 neural channels.

### Top Performing Neural Decoder

Given the audio data set of 5 English words and using all 96 channels, the top performing neural decoder, as ranked by validation set ESTOI, had the properties presented in Table 2.

**Table 2:**
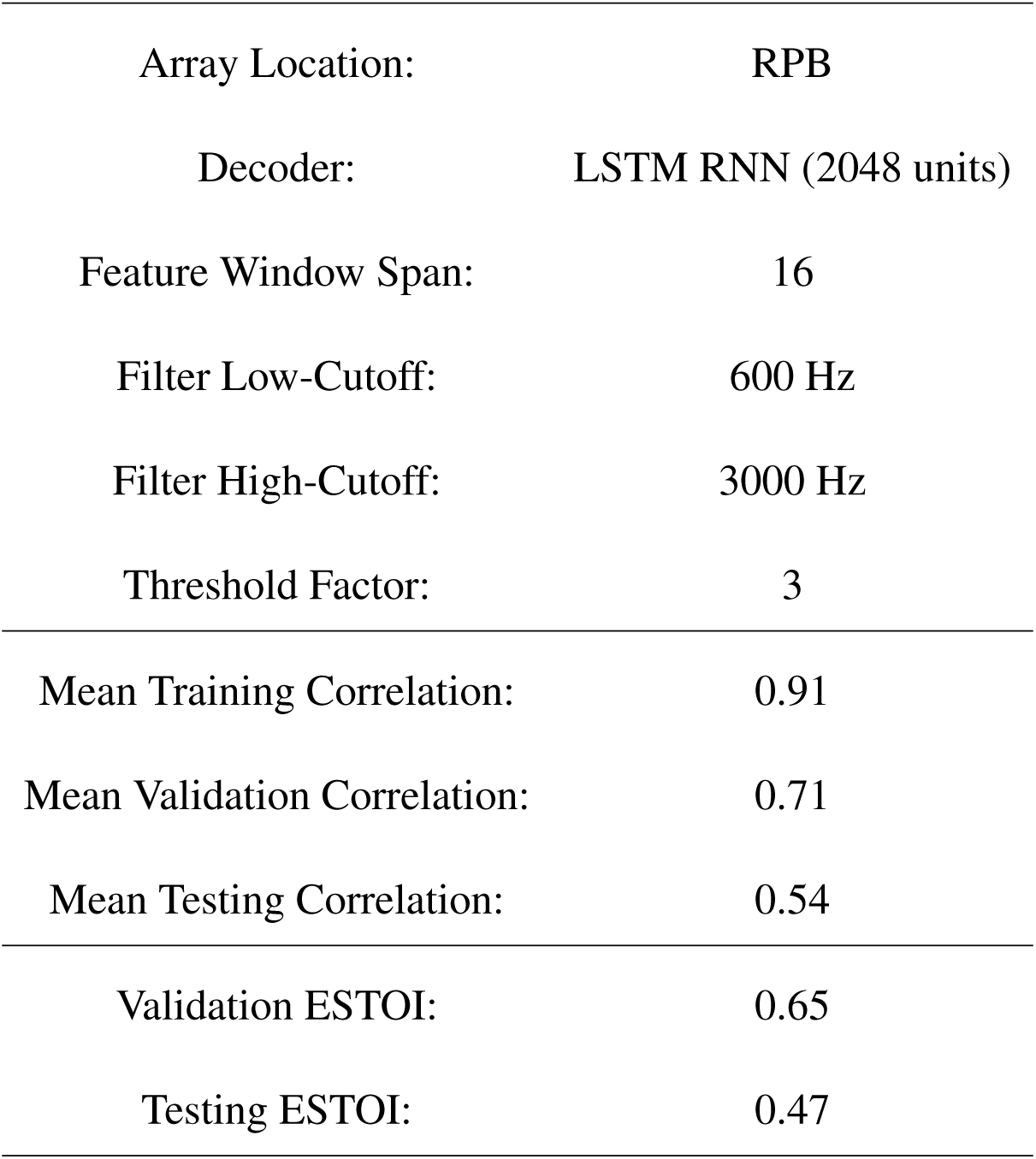
The hyperparameters and performance of the top performing neural decoding model that successfully reconstructed audio heard by a listener from multichannel neural spiking data recorded in STG.

This model successfully reconstructed intelligible audio from STG neural spiking activity with a validation ESTOI of 0.65 and a testing ESTOI of 0.47.

## Discussion

The work presented in this paper builds on prior foundational work in non-human primates and human subjects in mapping and interpreting the role of the secondary auditory cortex by intracranial recording of responses to external auditory stimuli. In this paper, we hypothesized that the STG of the secondary auditory cortex is part of a powerful cortical computational network for processing sounds. In particular, we explored how complex sounds such as English language words were encoded even if unlikely to be cognitively recognized by the macaque subject.

We combined two sets of methods as part of our larger motivation to take steps towards human cortical and speech prosthesis while leveraging the accessibility of the macaque model to chronic electrophysiology. First, use of 96-channel intracortical MEAs was speculated to yield a possibly important new view of STG neural activity via direct recording of neural population (spiking) dynamics. To our knowledge, such multichannel implants have not been implemented in a macaque STG except for work that focused on the primary A1 and, to a lesser extent, the STG in this animal model^31^. Second, given the rapidly expanding use of deep-learning techniques in neuroscience, we sought to create a suite of neural processing tools for high speed parallel processing of neural decoding models. We also deployed methods developed for voice recognition and speech synthesis which are ubiquitous in modern consumer electronic applications. We demonstrated the reconstructed audio recordings successfully recovered the sounds presented to the NHP (see supplementary Summary Video^22^ and Correlation Video^25^).

We performed an end-to-end neural decoding grid-search to explore the effects of signal properties, algorithms, and hyperparameters on reconstructing audio from full-broadband neural data recorded in STG. This computational experiment resulted in neural decoding models that successfully decoded neural activity into intelligible audio on the training, validation, and test data sets. In this work, we found that the LSTM RNN decoder outperformed other common neural decoding algorithms. This is consistent with the findings of other studies in decoding neural activity recorded from the human motor cortex^32^, and recent work from our group has demonstrated the feasibility of using LSTM RNN decoders for real-time neural decoding^33^. We also showed that optimizing the neural frequency content via a bandpass filter prior to multiunit spike extraction provided a marginal decoding performance improvement. One question is whether future auditory neural prostheses may achieve practically useful decoding performance while sampling at only a few kilohertz.

We observed no consistent effect when varying the threshold factor used in extracting multiunit spike counts. When using an LSTM RNN neural decoder, longer windows of data and more LSTM nodes in the network improved decoding performance. In general, we found the complexity of the decoding task to affect the optimal channel count with increased channel counts improving performance on more complex audio data sets. However, decoders overfit the training data when the number of neural channels was too high relative to the audio complexity (i.e. number of sounds). Due to these results, we will continue to grid-search threshold factor, LSTM network size, and the number of utilized neural channels during future experiments.

While we observed some improvement in decoding performance from the MEA implanted in the rostral parabelt STG compared to the array implanted in the caudal parabelt STG, this work does not fully answer the question whether location specificity of the MEA is critical within the overall cortical implant area, or whether reading out from the spatially extended cortical network in the STG is relatively agnostic to the precise location of the multichannel probe. Nonetheless, we plan to utilize information simultaneously recorded from two or more arrays in future work to test if this enables the decoding of increasingly complex audio (e.g. English sentence audio or sequences of macaque calls recorded in the home colony). We are developing an active listening task for future experiments that will allow the NHP subject to directly report different auditory percepts. Note that reasonable sound reconstruction could be achieved with relatively few channels (see Figure 7). This is perhaps due to the relatively low complexity of the sounds and their rather short temporal duration. We note how the question of necessary and optimal neural information (channel counts) versus task complexity has been generally predicted theoretically^34^.

We also found that the audio representation and processing pipeline are critical to generating intelligible reconstructed audio from neural activity. Other studies have shown benefits from using deep-learning to enable the audio representation/reconstruction^21^, and future work will explore these methods in place of mel-compression/Griffin-Lim algorithm. While we will continue to improve the performance of our neural decoding models, the presented results provide one starting point for future neural encoding work to “write in” neural information by patterned microstimulation^35, 36^ to elicit naturalistic audio sensations. Such future work can leverage the results presented here in guiding steps towards the potential development of an auditory cortical prosthesis.

## Methods

### Research Subjects

This work included one male adult rhesus macaque. The animal had two penetrating MEAs (Blackrock Microsystems, LLC, Salt Lake City, UT) implanted in STG each providing 96 channels of broadband neural recordings. A second animal was also implanted and trained; however, that data is not presented here as it will be used in future work. All research protocols were approved and monitored by Brown University Institutional Animal Care and Use Committee, and all research was performed in accordance with relevant NIH guidelines and regulations.

### Brain Maps and Surgery

The rostral and caudal parabelt regions of STG have been shown to play a role in auditory perception^3, 37, 38^. Those two parabelt areas are closely connected to the anterior lateral belt and medial lateral belt which show selectivity for the meaning of sound (“what”) and the location of the sound source (“where”), respectively^4^. Here, our emphasis was to ask if and how multichannel intracortical population recordings might depend on the array location within the parabelt.

Our institutional experience with implanting MEAs in NHPs has suggested that there is a non-trivial failure rate for MEA titanium pedestals^39^. To enhance the longevity of the recording environment, we staged our surgical procedure in two steps. First, we created a custom planar titanium mesh designed to fit the curvature of the skull. This mesh was designed using a 3D-printed skull model of the target area (acquired by MRI and CT imaging) and was coated with Hydroxyapatite to accelerate osseointegration. This mesh was initially implanted and affixed with multiple screws, providing a greater surface area for osseointegration on the NHP’s skull. Postsurgical CT and MRI scans were combined to generate a 3D model showing the location of the mesh in relation to the skull and brain.

Several weeks after the first mesh implantation procedure, we devised a surgical technique to access the parabelt region. A bicoronal incision of the skin was performed. The incision was carried down to the level of the zygomatic arch on the left and the superior temporal line on the right. The rhesus macaque has a significant amount of temporal muscle which prevented an inferior temporal craniotomy. To provide us with lower access on the skull base, we split the temporal muscle in the plane perpendicular to our incision (i.e. creating two partial thickness muscle layers) which were then retracted anteriorly and posteriorly. This allowed us to have sufficient inferior bony exposure to plan a craniotomy over the middle temporal gyrus and the Sylvian fissure.

The mesh, lateral sulcus, superior temporal sulcus, and central sulcus served as reference locations to guide the MEA insertion (see Figure 11). MEA arrays were implanted with a pneumatic inserter^39^.

**Figure 11:**
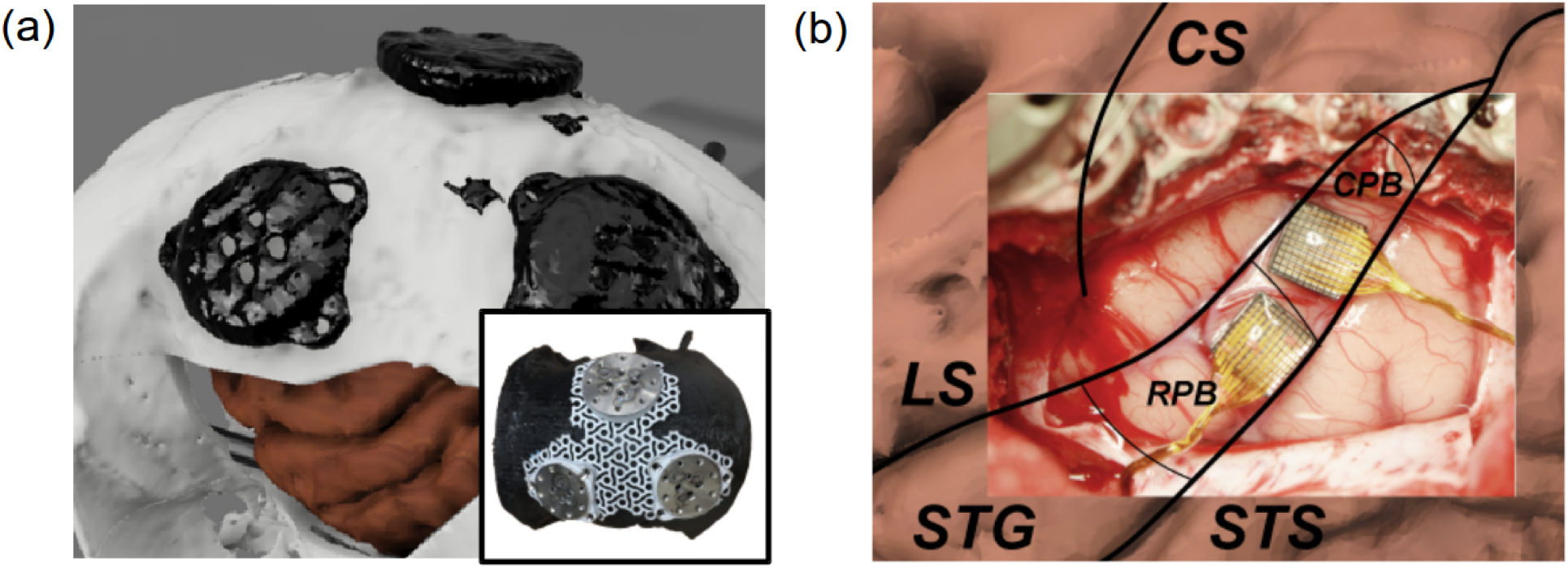
(a) A 3D model of the skull, brain, and anchoring metal footplates (constructed by merging MRI and CT imaging). Also, a titanium mesh on a 3D printed skull model. (b) Photo of the exposed area of the auditory cortex with labels added for relevant cortical structures (CS: central sulcus, LS: lateral sulcus, STG: superior temporal gyrus, STS: superior temporal sulcus, RPB: rostral parabelt, CPB: caudal parabelt).

### NHP Training and Sound Stimulus

Broadband neural data were collected to characterize the mesoscale auditory processing of STG during passive listening. While an NHP was restrained in a custom chair located in an echo proof chamber, complex sounds (English words and a macaque call) were presented through a focal loudspeaker near the animal’s head. The subject was trained to remain stable during the training session to minimize audio artifacts.

The passive listening task was controlled with a PC running MATLAB (Mathworks Inc., Natick, MA, Version 2018a). Animals were rewarded after every session using a treat reward. Within one session, 30 stimuli representations were played at approximately 1 second intervals. Data from 5 to 6 sessions in total were collected in one day. Computer synthesized English words were chosen to have different lengths (1 and 2 syllables) and distinct spectral contents (see Figure 12).

**Figure 12:**
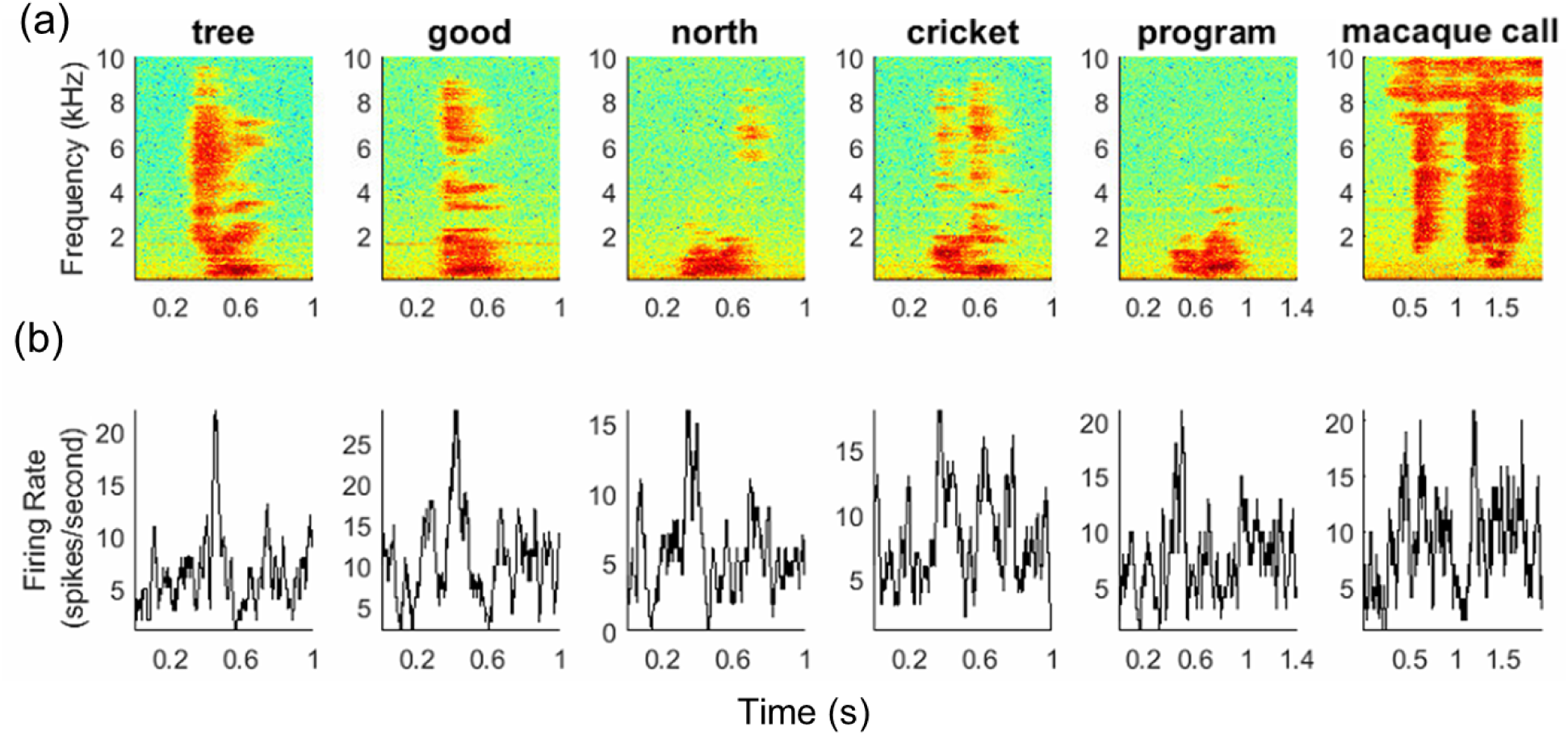
Neural data from the RPB array. (a) Spectrograms of five English word sounds and one macaque call. (b) Histogram for multiunit spiking activity on a given recording channel. In all plots, the spectrogram hop size is 2 ms and the window size for firing rates is 24 ms. Note the different scales of the horizontal axes due to different sound lengths.

In this work, a total of 5 different words were chosen and synthesized using MATLAB. Each sound was played 40-60 times with the word presentations pseudorandomly ordered. Trials that showed audio artifacts caused by NHP movement or environmental noise were manually rejected before processing.

### Intracortical Multichannel Neural Recordings

We used MEAs from Blackrock Microsystems with an iridium oxide electrode tip coating that provided a mean electrode impedance of around 50 kOhm. Iridium oxide was chosen with future intracortical microstimulation in mind. The MEA electrode length was 1.5 mm and 0.4 mm pitch for high-density grid recording. Intracortical signals were streamed wirelessly at 30 kS/s per channel (12-bit precision, DC) using a CerePlex W wireless recording device^23, 24^. Primary data acquisition was performed with a Digital Hub and Neural Signal Processor (NSP) (Blackrock Microsystems, Salt Lake City, UT). Audio was recorded at 30 kS/s synchronously with the neural data using a microphone connected to an NSP auxiliary analog input. The NSP broadcasted the data to a desktop computer over a local area network for long-term data storage using the Blackrock Central Software Suite. Importantly, the synchronous recording of neural and audio data by a single machine guaranteed that the neural data and audio data were aligned for offline neural decoding/encoding machine learning analysis.

### Neural Processing Toolkit and Cloud Infrastructure

We developed a neural processing toolkit (NPT) for performing large-scale distributed neural processing experiments in the cloud. The NPT is fully compatible with the ExOS machine learning operating system (Connexon Systems, Providence, RI). This enabled us to integrate software modules coded in different programming languages to implement processing pipelines. In total, the presented neural decoding analysis was composed of 26 NPT software modules coded in the Python, MATLAB, and C++ programming languages.

ExOS was used to launch, manage, and monitor experiments run on Google Cloud Platform (GCP) Compute Engine virtual machines (VMs). For this work, experiments were dispatched to a cluster of 10 VMs providing a total of 960 virtual CPU cores and over 6.5 terabytes of memory. Future experiments will leverage GPUs for accelerating deep-learning. Experiment configuration files implemented the described grid-search by instantiating NPT modules with different combinations of parameters and defining dependencies between modules for synchronized execution and data passing. ExOS automatically extracted concurrency from these experiment configurations thereby scaling NPT execution across all available cluster resources. The presented neural processing experiments evaluated 6,059 unique neural decoding models.

### Neural Preprocessing

We used multiunit spike counts (see Figure 12) as neural features for the neural decoders. We leveraged the Combinato^40^ library to perform spike extraction via a threshold crossing technique. Raw neural data was first scaled to units of microvolts and filtered using a 2nd-order bandpass elliptic filter with grid-searched cutoff frequencies. A noise level (i.e. standard deviation) of each channel was estimated over the training set using the median absolute deviation to minimize the interference of spikes^27^. We set thresholds for each individual channel by multiplying the noise levels by a threshold factor. We detected negative threshold crossings on the full-broadband 30 kS/s data and then binned them into counts.

### Audio Preprocessing

We manually labeled raw audio to designate the begin and end time indices for single sound trials. The audio data contained 5 different English words as well as a single macaque call. We subselected sounds to create target audio data sets for the neural decoders with varying complexity. The librosa audio analysis library^41^ was used to calculate the short-time Fourier transform spectrogram with an FFT window of 2048. This spectrogram was then compressed to its mel-scaled spectrogram to reduce its dimensionality to the number of melbands. These mel-bands served as the target data for the evaluated neural decoders. Audio files were recovered from the mel-bands by inverting the mel-spectrogram and using the Griffin-Lim algorithm^28^ to recover phase information. All targets were standardized to zero-mean using scikitlearn^42^ transformers fit on the training data.

### Decoding Algorithms

We evaluated 7 different neural decoding algorithms based on the KordingLab Neural Decoding library^43^ including a Kalman filter, Wiener filter, Wiener cascade, Dense neural network, simple recurrent RNN, GRU RNN, and LSTM RNN. Each neural network consisted of a single hidden layer and and an output layer. Hidden units in the dense neural network and simple RNN used rectified linear unit activations^44^. The GRU RNN and LSTM RNN used tanh activations for hidden units. All output layers used linear activations, and no dropout^45^ was used. We used the adam optimization algorithm^46^ to train the dense neural network and RMSprop^47^ for all recurrent neural neural networks. We sequentially split the data set (~40 trials per sound per array sampled in 40ms bins) into training, validation, and testing sets composed of 80%, 10%, and 10% of the data, respectively. We trained all neural networks using early stopping^48^ with a maximum of 2048 training epochs and a patience of 5 epochs. Mean-squared error was used as the monitored loss metric.

### Code Availability

The neural processing toolkit code used in this study is available upon request from the corresponding author [C.H.].

### Data Availability

The data that supports the findings of this study are available upon request from the corresponding authors [C.H., J.L., A.N.].

## Supporting information

Summary Video

Correlation Video

## Acknowledgements

The authors express their gratitude to Laurie Lynch for her expertise and leadership with our macaques. We also thank Huy Cu, Carlos Vargas-Irwin, Kevin Huang, and the Animal Care facility at Brown University for their most important contributions. We thank Rosalind Mandelbaum for her contributions to the supplementary videos. We are most appreciative to Josef Rauschecker for generously sharing his deep insights into the role and function of the primate auditory cortex.

This research was initially supported by Defense Advanced Research Projects Agency N66001-17-C-4013, with subsequent support from private gifts.

## Author Contributions

A.N. and J.L. conceived the project. J.L. and A.N. designed the neural experimental concept. J.L. designed and executed the NHP experiments. C.H. designed and executed the neural decoding machine learning experiments. C.H. and R.O. developed the neural processing toolkit. C.H. performed the results analysis. W.T. provided the neurocomputational expertise and D.B. the surgical leadership. C.H., A.N, and J.L. wrote the manuscript. All authors commented on the manuscript. C.H. composed the supplementary videos.

